# Azithromycin resistance through interspecific acquisition of an epistasis dependent efflux pump component and transcriptional regulator in *Neisseria gonorrhoeae*

**DOI:** 10.1101/309294

**Authors:** Crista B. Wadsworth, Brian J. Arnold, Mohamad R. Abdul Sater, Yonatan H. Grad

## Abstract

Mosaic interspecifically acquired alleles of the multiple transferable resistance (*mtr*) efflux pump operon correlate with reduced susceptibility to azithromycin in *Neisseria gonorrhoeae* in epidemiological studies. However, whether and how these alleles cause resistance is unclear. Here, we use population genomics, transformations, and transcriptional analyses to dissect the relationship between variant *mtr* alleles and azithromycin resistance. We find that the locus encompassing the *mtrR* transcriptional repressor and the *mtrCDE* pump is a hotspot of interspecific recombination introducing alleles from *N. meningitidis* and *N. lactamica* into *N. gonorrhoeae*, with multiple rare haplotypes in linkage disequilibrium at *mtrD* and the *mtr* promoter region. Transformations demonstrated that resistance is mediated through epistasis between these two loci and that the full length of the mosaic *mtrD* allele is required. Gene expression profiling revealed the mechanism of resistance in mosaics couples the novel *mtrD*alleles with promoter mutations enhancing expression of the pump. Overall, our results demonstrate that epistatic interactions at *mtr* gained from multiple *Neisseria* has contributed to azithromycin resistance in the gonococcal population.

**AUTHOR SUMMARY:** *Neisseria gonorrhoeae* is the sexually transmitted bacterial pathogen responsible for over 100 million cases of gonorrhea worldwide each year. The incidence of reduced susceptibility to the macrolide class antibiotic azithromycin has increased in the past decade; however, a large proportion of the genetic basis of resistance to this drug remains unexplained. Recently, resistance has been shown to be highly associated with mosaic alleles of the multiple transferable resistance (*mtr*) efflux pump, which have been gained via horizontal gene exchange with other *Neisseria*. However, if and how these alleles caused resistance was unknown. Here, we demonstrate that resistance has been gained through epistasis between *mtrD* and the *mtr* promoter region using evidence from both population genomics and experimental genetic manipulation. Epistasis also acts within the *mtrD* locus alone, requiring the full length of the gene for phenotypic resistance. Transcriptomic profiling indicates that the mechanism of resistance in mosaics is likely derived from both structural changes to *mtrD*, coupled with promoter mutations that result in regulatory changes to *mtrCDE*.

## INTRODUCTION

The causal agent of gonorrhea, *Neisseria gonorrhoeae*, is a gram-negative diplococcus and an exclusively human pathogen. The prevalence of *N. gonorrhoeae* with reduced susceptibility to azithromycin has dramatically increased in recent years from just 0.6% in 2013 to 3.6% in 2016 in the United States [1], 0.8% in 2013 to 4.7% in 2016 in England and Wales [2], and 5.4% in 2013 to 7.1% in 2015 across Europe [3]. Additionally, reports in 2015 from both China and Japan have documented resistance in as high as 30% of the gonococcal population in some regions [4]. This spike in resistance is alarming, as azithromycin is one of the two first line drugs recommended as dual antimicrobial therapy for uncomplicated cases of gonococcal urethritis by the Centers for Disease Control (CDC). Azithromycin is a macrolide antibiotic that inhibits protein synthesis by binding to the 23s rRNA component of the 50S ribosome, and while the majority of resistance can be explained by mutations in the 23s rRNA azithromycin binding sites (i.e., C2611T and A2059G) [5–7] the genetic basis of ~36% of resistance is still unexplained within the U.S. population [7], thus limiting the potential for development of molecular-based resistance diagnostics and restricting our understanding of the evolutionary paths to reduced drug susceptibility.

Gonococci are adept at acquiring antimicrobial resistance as a result of their natural competence for transformation, allowing for the spread of resistance and other adaptively advantageous alleles, between lineages and even across species boundaries [8–11]. Extensive intragenus gene exchange has led to the concept of *Neisseria* as a consortium of species interconnected by allele sharing, with ‘fuzzy’ borders permitting rapid access to new adaptive solutions [11,12]. In gonococci, intragenus recombination is an important source of novel genetic variation with many observations of ‘mosaic’ loci gained from other neisserial species [13–16]. However, aside from horizontal gene transfer facilitating the evolution of resistance to third-generation cephalosporins through acquisition of mosaic *penA* [7,14], allelic exchange has not yet been proven to be the basis for resistance to any other antibiotic class in this species.

Recent epidemiological studies from the Unites States, Canada, and Australia have reported an association between mosaic *multiple transferable resistance* (*mtr*) efflux pump alleles and reduced susceptibility to azithromycin [7,17–19]. *Mtr* mosaics appear to have originated through horizontal gene exchange from other *Neisseria*, and have been identified by high sequence homology of the repressor of the pump (*mtrR*) to *N. meningitidis* and *N. lactamica*. Mosaics have previously been associated with an outbreak of azithromycin resistance in Kansas City, MO from 1999–2000 [20,21], and also the majority of azithromycin resistance reported in New South Wales, Australia [19]. While correlation between mosaic *mtr* and azithromycin resistance suggests causality, there is little experimental evidence to confirm the association.

The Mtr efflux pump is comprised of the MtrC-MtrD-MtrE cell envelope proteins, which together export diverse hydrophobic antimicrobial agents such as such as antibiotics, nonionic detergents, antibacterial peptides, bile salts, and gonadal steroidal hormones from the cell [22–26]. Mtr-mediated resistance to diverse antimicrobial agents in gonococcus is thought to act via enhanced drug export as a result of overexpression of *mtrCDE*. Mutations that alter expression of the pump include the *mtrC_120_* substitution, an adenine to guanine transition located 120 bp upstream of the *mtrC* start codon which acts as an alternative promoter for *mtrCDE* [27,28]; an A-deletion in the *mtrCDE* promoter that has been shown to repress the transcription of *mtrR* while simultaneously enhancing transcription of *mtrCDE* [29]; and mutations that abrogate the function of MtrR by inducing premature stop codons or radical amino acid substitutions in the DNA-binding motif [20,30,31]. However, it is unclear if resistance in *mtr* mosaics is derived from any of these mechanisms.

Here, we used a combination of population genomic and experimental approaches to dissect the mechanism of resistance in mosaics. We first assessed patterns of allelic diversity within the gonococcal population to define the boundaries of horizontal gene transfer at *mtrRCDE*, and found that the entire *mtr* region is a hotspot of interspecies recombination which has introduced multiple rare and divergent mosaic alleles from *N. meningitidis* and *N. lactamica* into the gonococcal population. Strong linkage disequilibrium at *mtrD* and the *mtr* promoter region suggested the maintenance of epistatic allelic combinations, thus we tested for interaction effects within and between *mtr* loci via transformation. We discovered epistatic interactions across almost the entirety of *mtrD*, and also between mosaic *mtrD* and mosaic *mtr* promoter regions, that synergistically enhanced azithromycin resistance. Furthermore, patterns of diversity in this region coupled with experimental evidence suggest antibiotic-mediated selection may be acting on these epistatic interactions. Finally, we tested for regulatory evolution of pump components, as previous mechanisms of azithromycin resistance through the Mtr efflux pump have been demonstrated to be expression-driven. Our results support that inheritance of mosaic promoter regions increases the expression of *mtrCDE* while gaining mosaic *mtrD* alone does not. Thus, the likely mechanism of resistance in mosaics is a structural change to *mtrD*, which enhances the capacity of the protein to recognize or transport azithromycin, coupled with increased efflux through the amplified production of pump components.

## RESULTS

### Allelic diversity suggests increased interspecies admixture at *mtrRCDE*

To gain insight into the evolutionary history of the *mtrR* transcriptional repressor and the *mtrCDE* pump, we analyzed patterns of diversity using the 1102 Gonococcal Isolate Surveillance Project (GISP) isolates described in Grad et al. [7,32]. A significant increase in allelic diversity was observed across *mtrRCDE* compared to the entire genome, with the highest diversity at *mtrD* (Figure 1a,b; Supplementary Table 1). We also detected a significant enrichment of rare alleles in the population across *mtrRCDE* (Figure 1a,c; Supplementary Table 1). Linkage disequilibrium was strongest at *mtrD* and the *mtr* promoter region in a comparison of all pairs of single nucleotide polymorphisms (SNPs) within *mtrRCDE*, with higher linkage observed at pairs of variant sites within each of these loci (Figure 1d,f; Supplementary Table 1).

**Figure 1.**
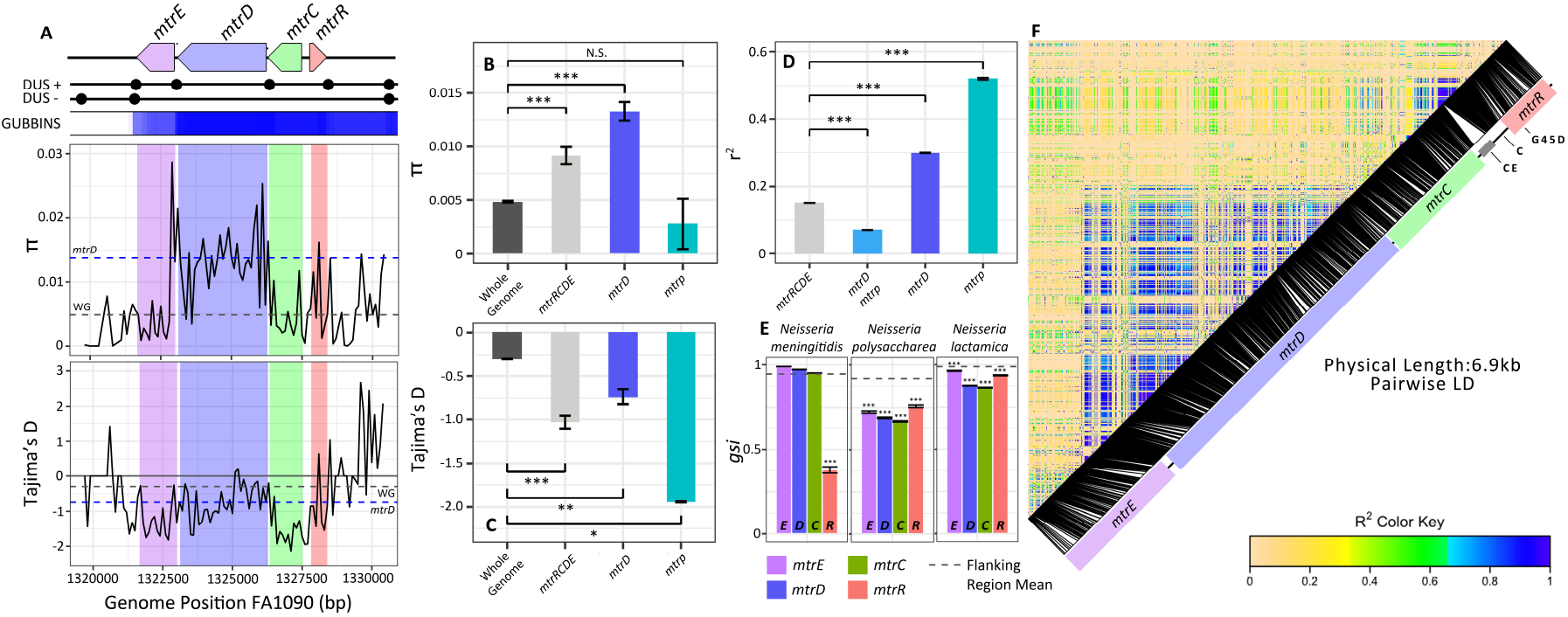
Horizontal gene transfer (HGT) of *mtr* introduces novel adaptive genetic variation into *Neisseria gonorrhoeae*. (A-C) The U.S. gonococcal population (n=1102 isolates) shows patterns of elevated allelic diversity across *mtrRCDE*, with the highest diversity at *mtrD*, compared to the rest of the genome. An excess of rare alleles across *mtrRCDE* (Tajima’s D<0) suggests the introduction of new genetic variation in this region has occurred recently, possibly after a genome-wide selective sweep or population contraction. (E) Depressed *gsi* values indicate importation of divergent alleles from multiple neisserial species into gonococcus across all *mtr* loci. (D,F) The strongest linkage was observed within *mtrD* and the *mtr* promoter regions.

**Table 1.** Properties of strains used in the study

| Strain | Source | AZI<sup>a</sup> MIC (µg/ml)* |
| --- | --- | --- |
| GCGS0276 | GISP<sup>b</sup> isolate, Kansas City | 1 |
| GCGS0402 | GISP<sup>b</sup> isolate, Miami | 2 |
| GCGS0834 | GISP<sup>b</sup> isolate, Los Angeles | 2 |
| GCGS0353 | GISP<sup>b</sup> isolate, Dallas | 0.03 |
| GCGS0465 | GISP<sup>b</sup> isolate, Phoenix | 0.06 |
| 28B1 | Disseminated gonococcal infection isolate (CDC, 1974) | 0.125 |

*a* AZI, azithromycin
*b* GISP, *Gonococcal Isolate Surveillance Project*
\*MIC scores are the average of three independent replicate tests.

To define interspecific admixture events within *Neisseria*, we characterized the genealogical sorting index (*gsi*; [33]) to explore gene tree topology measures of species-specific phylogenetic exclusivity. *gsi* ranges from 0 (no exclusivity) to 1 (monophyletic), and serves as a metric to assess allele sharing that may arise through interspecific recombination or incomplete divergence from a recent split between species. We calculated *gsi* for genes in a 50 kb window encompassing *mtrRCDE* with conserved microsynteny between *N. gonorrohoeae* and other *Neisseria* (Supplementary Figure 1). This region included 29 core genes excluding *mtrRCDE*. To define the region-specific *gsi* background, we calculated *gsi* values across 100 bootstrap replicates for each gene in the *mtrRCDE* flanking region by species. Mean gonococcal *gsi* was 0.95 with *N. meningiditis*, 0.99 with *N. lactamica*, and 0.92 with *N. polysaccharea* for flanking genes (Figure 1e; Supplementary Table 2). Significant reductions in *gsi* were detected across *mtrRCDE* compared to the 29 loci within the surrounding 50 kb region (Figure 1e). Significant allele sharing between *N. gonorrhoeae* and *N. meningiditis* was exclusively observed at *mtrR* (Figure 1e; Supplementary Table 2), while significant allele sharing between *N. gonorrhoeae* and both *N. lactamica* and *N. polysaccharea* occurred across all *mtr* loci (Figure 1e; Supplementary Table 2).

There were multiple recombined mosaic haplotypes present spanning the full-length of *mtrD* (n=80), *mtrRCD* (n=9), *mtrRCDE* (n=20), and some isolates with partial mosaic *mtrD* with the majority of the gene homologous to native gonococcal sequence (n=13) (Figure 2). Of the 109 isolates with full-length mosaic *mtrD*, 4 were 99% identical to *N. meningitidis*, 5 had alleles with 94–96% identity to *N. lactamica*, and the remainder had alleles that aligned equally well to *N. meningitidis* and *N. lactamica* with identities ranging from 91–92%. Of the 29 isolates with mosaic promoter regions identified by Grad et al. [7], 24 were 96–98% identical to *N. lactamica*, 4 were 99% identical to *N. meningitidis* with the presence of a 153-bp Correia element insertion [20], and 1 was 92% similar to *N. meningitidis* but lacked the Correia element that was present in the other four *N. meningitidis*-like isolates. All isolates with full-length mosaic *mtrD* had azithromycin MICs ≥ 0.25 μg/ml, while all isolates with full-length mosaic *mtrD* and a mosaic *mtr* promoter had MICs ≥ 1 μg/ml (Figure 2).

**Figure 2.**
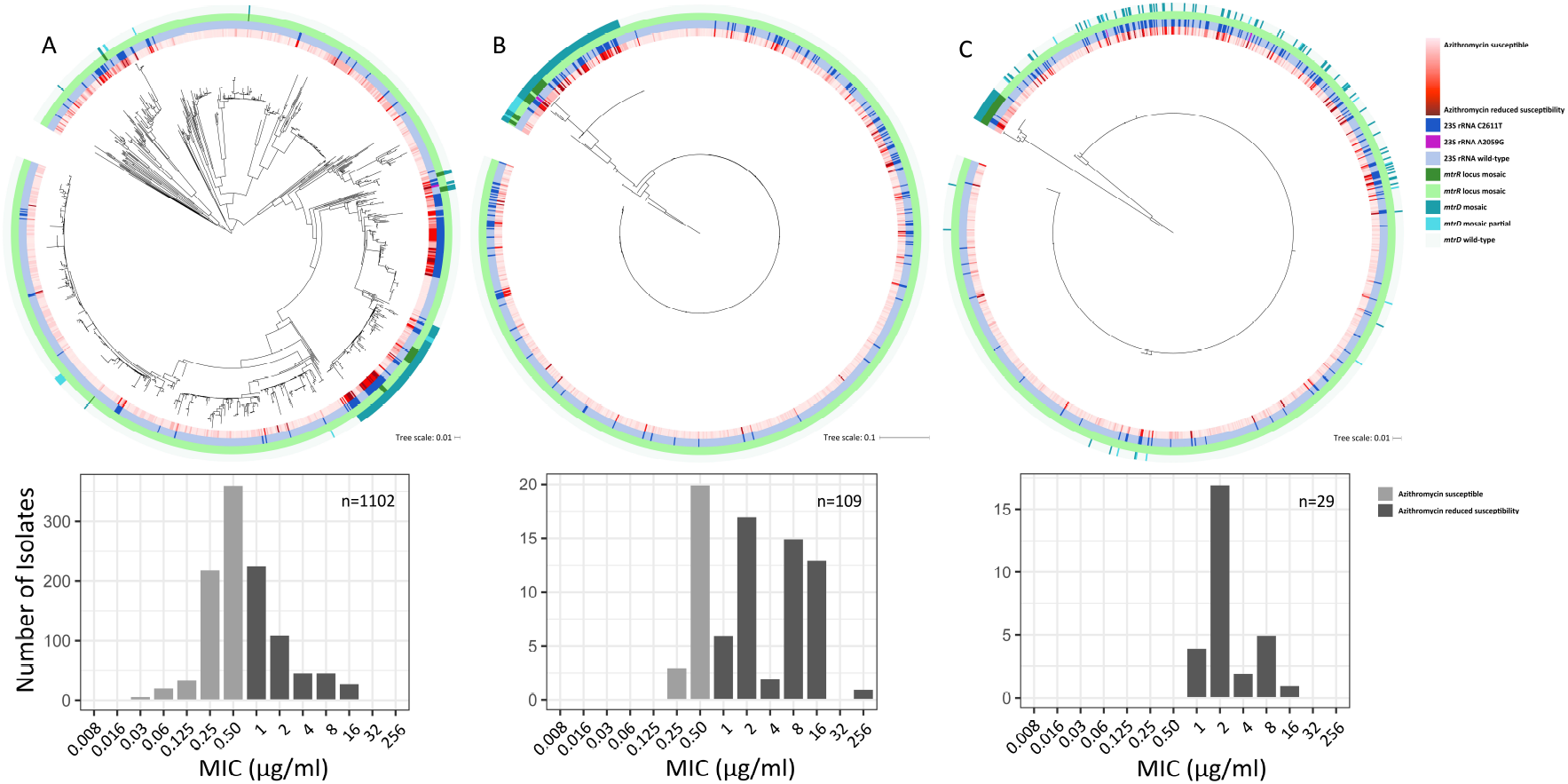
Divergent *mtrD* and *mtr* promoter haplotypes are associated with step-wise increases in MIC to azithromycin. (A) A maximum likelihood whole-genome-sequence phylogeny of 1102 *Neisseria gonorrhoeae* isolates, based on single-nucleotide polymorphisms generated from mapping to the FA1090 reference genome (Grad et al. 2016), is associated with a distribution of MIC values which fall both above and below the defined resistance threshold (MIC > 1 μg/ml). The inner annotation ring shows MICs to azithromycin on a continuous scale, the following annotation ring indicates isolates with at least 2 copies of the C2611T 23S ribosomal RNA (rRNA) mutation or isolates with 4 copies of the A2059G 23S rRNA mutation, the next annotation ring shows isolates that were identified as interspecies mosaics based on their sequence at *mtrR* by Grad et al. (2016), and the outermost annotation ring shows isolates identified as *mtrD* mosaics in this study. (B) A maximum likelihood phylogeny built on *mtrD* alignments show 109 isolates with full-length mosaic alleles at this locus associated with elevated MICs to azithromycin. (C) A maximum likelihood phylogeny built on the *mtr* promoter region identifies all 29 mosaics with reduced susceptibility to azithromycin identified by Grad et al. [7] also have inherited mosaic *mtrD*.

**Table 2.**
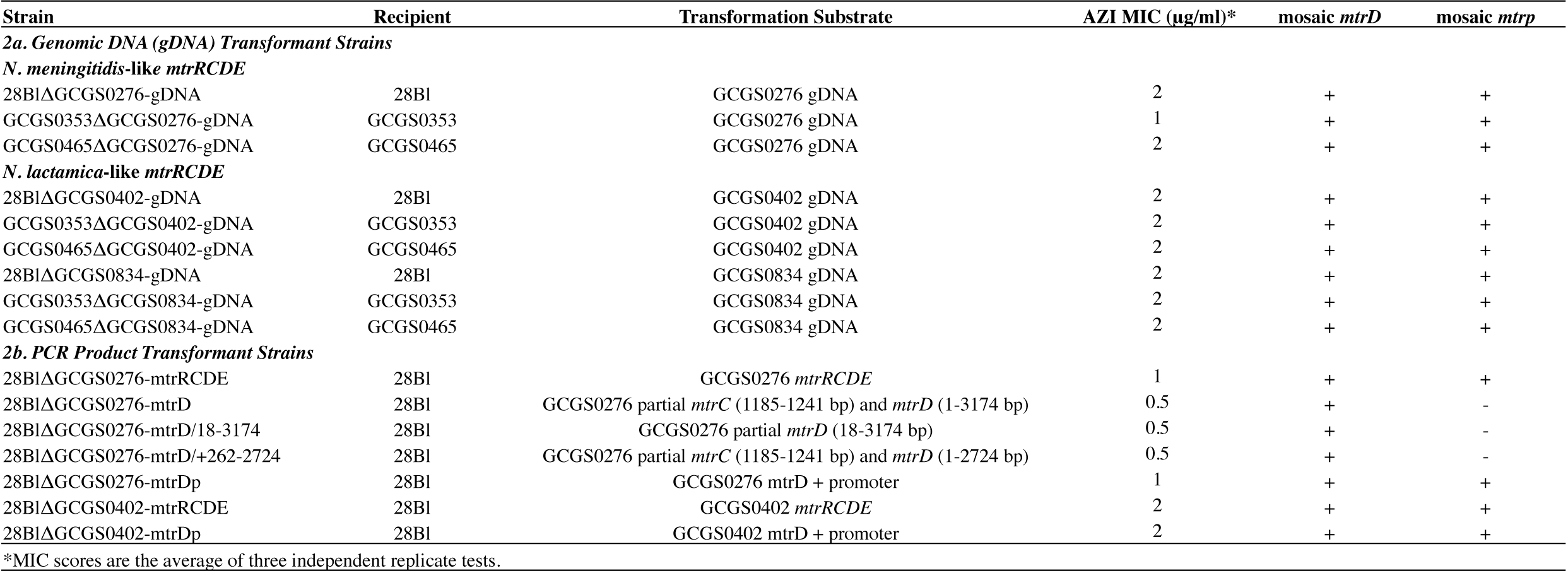
MIC values of transformant strains

Of the twenty nine *mtr* mosaics described in Grad et al. [7], none had the *mtrC_120_* substitution, A-deletion, 23s ribosomal rRNA mutation A2059G, mutations in *rplD, rplV* tandem duplications, or variants of the rRNA methylases *ermC* and *ermB* that have been associated with or experimentally confirmed to be involved in azithromycin resistance [6,7,18,23,28,29,34]. However, four isolates had the premature stop codon mutations in *mtrR*, and five had the C2611T 23s rRNA mutation [5].

### Epistasis between multiple *mtr* loci and within *mtrD*

We exploited the natural competence of *Neisseria* to explore the potential for mosaic *mtr* alleles to produce reduced susceptibility to azithromycin by transforming susceptible strains with either genomic DNA (gDNA) or PCR-amplified products from mosaic donors. Susceptible recipient strains for transformations included: 28Bl [20,35,36], GCGS0353, and GCGS0465 (MIC ≤ 0.125 μg/ml; Table 1). Three strains with reported mosaic *mtr* alleles and azithromycin minimum inhibitory concentrations (MICs) ≥ 1 μg/ml were selected as donors for DNA transfer (Table 1). These isolates included GCGS0276, GCGS0834, and GCGS0402. Of these mosaics, GCGS0276 had a *N. meningitidis-like mtrR* sequence, while GCGS0834 and GCGS0402 had *N. lactamica-like mtrRs*. None of the donor strains had premature stop codons in *mtrR* or the C2611T mutation.

Genomic DNA from GCGS0276, GCGS0402, and GCGS0834 transformed multiple susceptible isolates to resistance (Table 2a). To identify the locus responsible, we sequenced the genomes of 28Bl cell lines transformed with gDNA from mosaic donors and characterized SNPs that had been inherited from donor strains that were not present in the 28Bl recipient. The only common region that had been inherited across all transformants was *mtrRCDE* (Supplementary Figure 2).

Genomics results indicated the presence of linkage disequilibrium at *mtrD* and the *mtr* promoter region (Figure 1 d,f). Thus, to test for possible interaction effects that contribute to antibiotic-dependent fitness, and to further characterize the mechanism underlying reduced susceptibility in *mtr* mosaics, we designed targeted amplicons for transformation from *N. meningitidis-like* mosaic (GCGS0276) and a *N. lactamica-like* mosaic (GCGS0402). For GCGS0276, the only locus within the *mtrRCDE* operon that was found to increase resistance to azithromycin alone was *mtrD* (Figure 3a; Table 2b). GCGS0276 *mtrD* in the 28Bl background raised the MIC to azithromycin by 3 fold, from 0.125 to 0.5 μg/ml, yet no single region of *mtrD* was able to produce the 0.5 μg/ml phenotype (Figure 3b). However, inheriting amplicons that contained both the 5’ (18–356 bp) and 3’ (2356–2724 bp) ends of GCGS0276 *mtrD* were together sufficient to increase the 28Bl MIC to 0.5 μg/ml (Figure 3c-e). There were four changes at the amino acid level between GCGS0276 and 28Bl in these regions, two in the PN1 domain of MtrD (I48T and G59D) and two in the PC2 domain (K823D and F854L) (Supplemental Figure 3); and in total twenty amino acid changes between the GCGS0276 and 28Bl proteins (Supplementary Figure 3). GCGS0402 *mtrD* alone was not able to produce resistance in 28Bl.

Transformants that inherited the entire *mtrRCDE* operons of GCGS0276 and GCGS0402 had MICs of 1 and 2 μg/ml respectively, mirroring the donor strain phenotypes. Thus, amplicons were designed for each of these strains to amplify the *mtrD* locus in combination with other regions of the operon to determine the combination of loci that would reproduce the donor resistance phenotypes. For both GCGS0276 and GCGS0402, we found that donor resistance phenotypes of 1 and 2 μg/ml could be produced in 28Bl by transforming both *mtrD* and the *mtr* promoter region together (Figure 4; Table 2b; Supplementary Figure 4).

**Figure 3.**
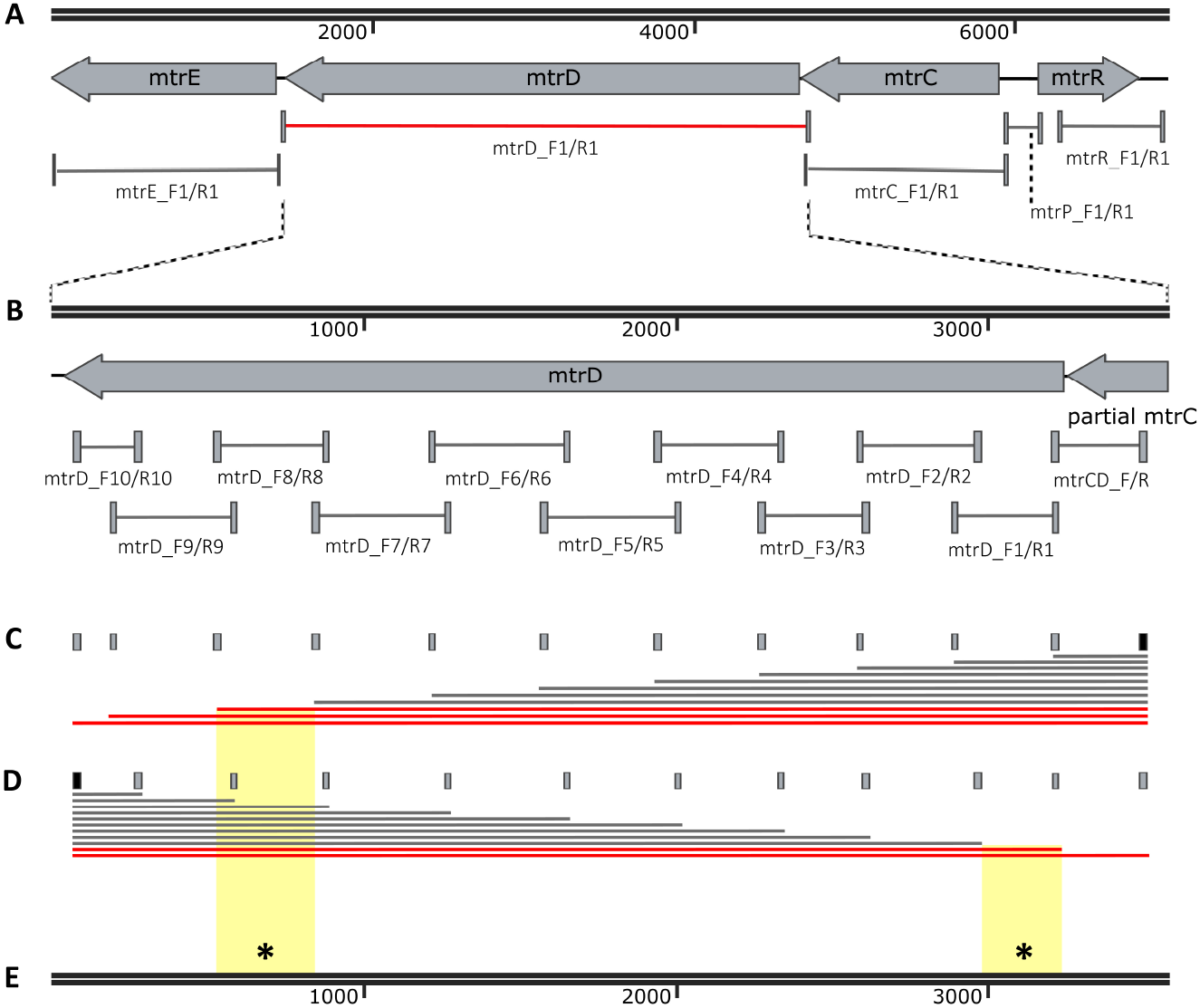
Epistatic interactions between multiple domains of *mtrD* contribute to elevated MICs. (A) GCGS0276 *mtrD* in the 28Bl background elevated MIC from 0.125 μg/ml to 0.5 μg/ml (red). (B) Primer pairs designed to amplify ~300 bp fragments over the length of *mtrD* resulted in no observed transformants on 0.38 μg/ml selection plates, suggesting multiple mutations across *mtrD* are needed for resistance. To determine the regions that contributed to resistance, multiple fragment sizes were constructed by (C) holding the rightmost forward primer (black) constant while adding different reverse primers (grey), and (D) holding the leftmost reverse primer (black) constant while adding forward primers (grey) to separate reactions. (E) At minimum SNPs at base pair positions 18 to 356 coupled with SNPs at positions 2356 to 2724 (*) were needed to raise the MIC from the 0.125 μg/ml of the recipient 28Bl strain to 0.5 μg/ml.

**Figure 4.**
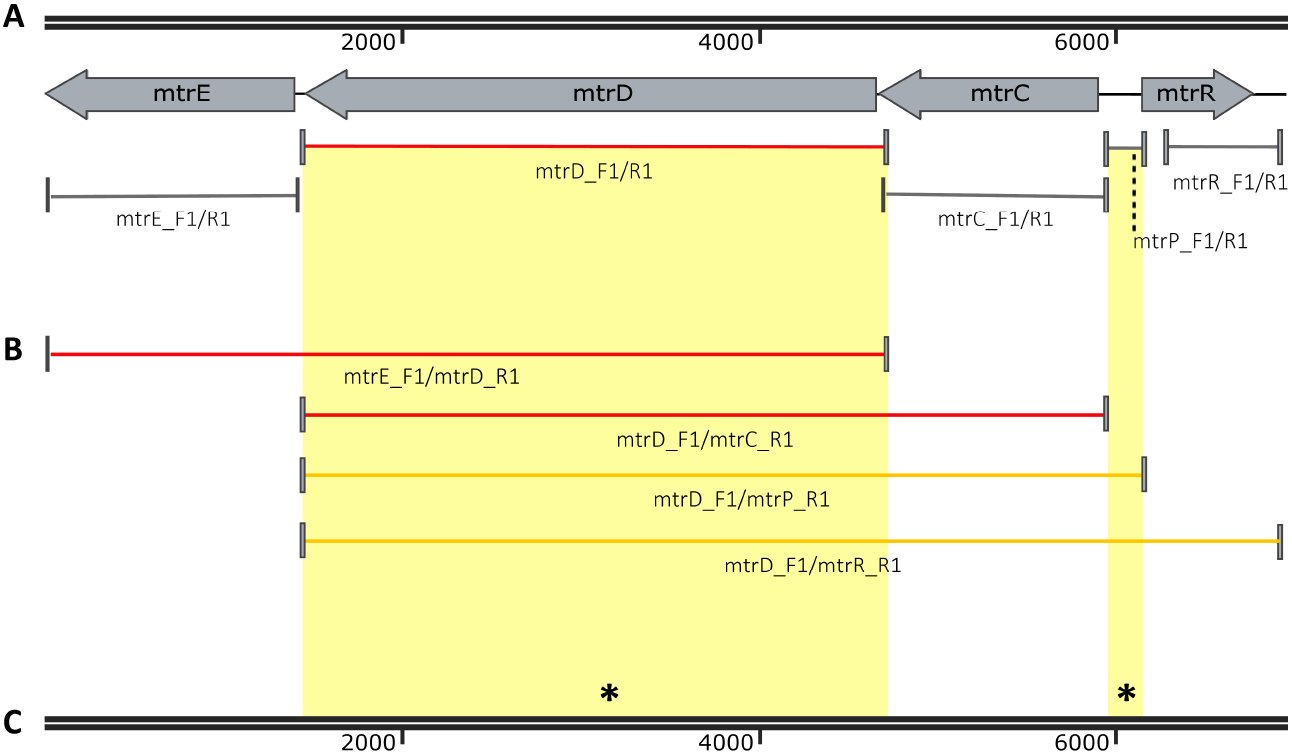
Epistasis between *mtrD* and the *mtr* promoter region is causal to reduced susceptibility. (A) GCGS0276 *mtrD* was the only region that could independently raise MIC from 0.125 to 0.5 μg/ml in the 28Bl background (red). (B) Only the addition of the *mtr* promoter raised MIC to the donor strain phenotype of 1 μg/ml (yellow lines).

### Regulatory and structural mutations epistatically contribute to resistance

We tested for the contribution of transcript regulatory variation to the mechanism of resistance by profiling gene expression via RNA-seq of 28Bl, 28BlΔGCGS0276-mtrD, and 28BlΔGCGS0276-mtrRCDE. As expression of the *mtr* efflux pump is inducible by exposure to antimicrobial agents [37,38], we evaluated expression pre-azithromycin exposure and 120 minutes after the addition of a sub-MIC dose of azithromycin (0.125 μg/ml) to the culture media. Across 24 libraries, a total of 106 million 50 bp paired-end reads mapped to the FA1090 reference genome. Each library had on average 4.44±3.49 million mappable reads.

We assessed the impact of mosaic *mtrD* on *mtrRCDE* mRNA expression by comparing 28BlΔGCGS0276-mtrD transformants to 28Bl, and found no significant differential regulation of transcripts encoding *mtr* efflux pump components (Supplementary Figure 5; Supplementary Table 3). To determine the effect of the mosaic *mtr* promoter on pump expression, we compared 28BlΔGCGS0276-mtrD and 28BlΔGCGS0276-mtrRCDE transformants (Figure 5; Supplementary Table 3). Here, presence of a mosaic *mtr* promoter resulted in the significant upregulation of *mtrC, mtrD*, and *mtrE* across conditions (FDR < 0.0001) and upregulation of *mtrR* in the absence of azithromycin (FDR=0.003).

**Figure 5.**
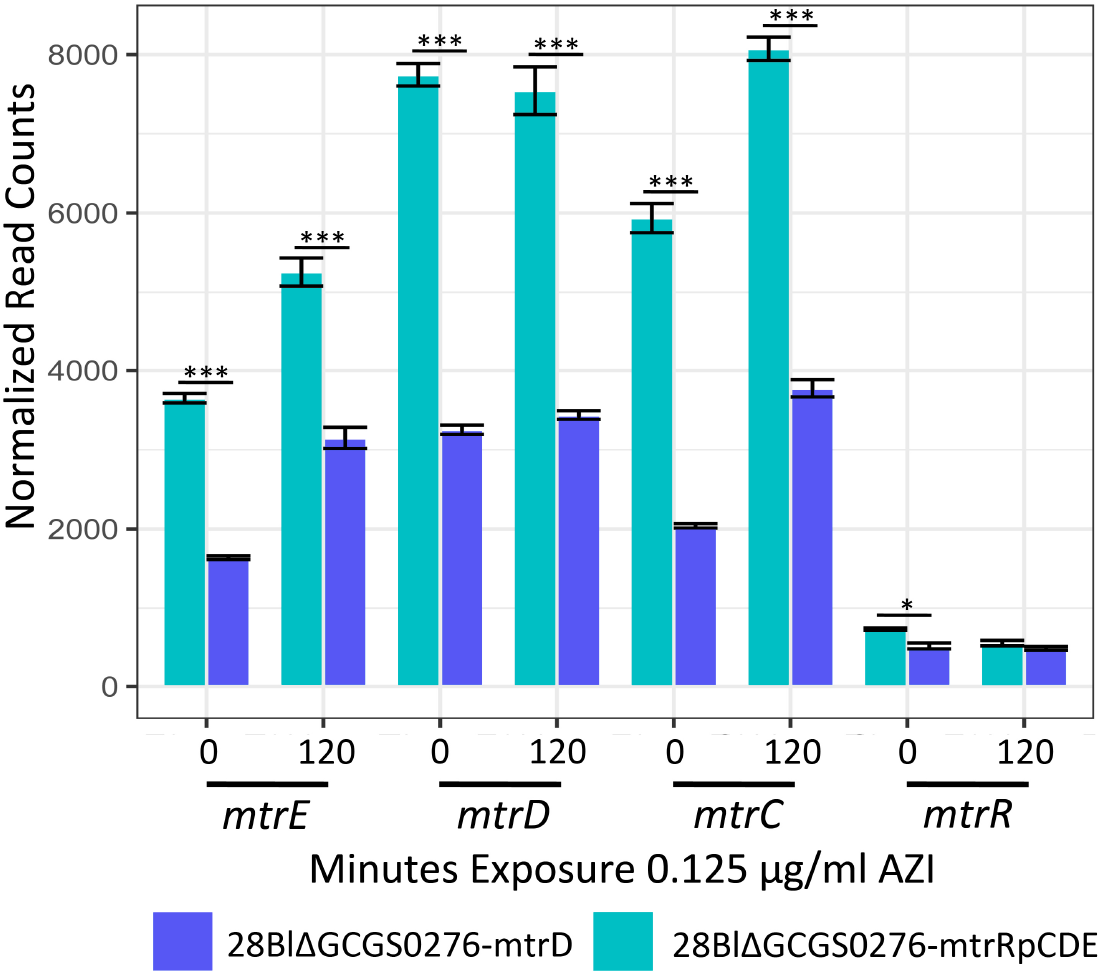
The *N. meningitidis*-like GCGS0276 mosaic *mtr* promoter sequence upregulates expression of *mtr* efflux pump component mRNAs. 28Bl transformants with mosaic GCGS0276 *mtrD* (blue) or mosaic GCGS0276 *mtrRCDE* (teal) were exposed to sub-MIC (0.125 μg/ml) concentrations of azithromycin for 120 minutes. In both the presence and absence of drug, the presence of the *mtr* promoter region results in significantly upregulated pump component mRNAs (FDR < 0.0001).

## DISCUSSION

Genomic and surveillance efforts from across the globe have demonstrated an association between mosaic *mtr* alleles and low-level azithromycin resistance [7,17–19], but the causal role of these alleles in generating resistance has been unclear. Mtr-mediated decreased susceptibility to macrolides has been attributed exclusively to increased *mtrCDE* expression [27–30]. However, a majority of *mtr* mosaic isolates in the population genomic dataset have none of the variants that alter regulation of *mtrCDE*, such as the *mtrC_120_* substitution, A-deletion, or premature stop codon mutations in *mtrR* [20,27–31].

Using a combination of experimental and population genomic approaches, we demonstrated that the mosaic *mtr* alleles are responsible for resistance and showed that the mechanism of resistance involves multiple loci, including an epistatic interaction between the *mtrD* component of the pump and the *mtr* promoter. Population genomics and phylogenetic reconstruction demonstrated at least twelve independent acquisitions of mosaic *mtr* alleles, which have introduced multiple rare *mtr* haplotypes from *N. meningitidis* and *N. lactamica* into the gonococcal population (Figure 1 and 2).

Despite the sequence divergence (8%) between *N. meningitidis* and *N. lactamica-like* mosaics, *mtr* sequences from both generated azithromycin resistance through the same mechanism. For the *N. meningitidis* mosaic, *mtrD* alone was able to raise the MIC of 28Bl to 0.5 μg/ml independent of transcriptional changes to the pump’s regulation; with the donor phenotype of 1 μg/ml only reproduced by adding the *N. meningitidis mtr* promoter region, which increased expression of *mtrCDE*. Similarly, although acquisition of the *N. lactamica mtrD* was not sufficient on its own to enhance resistance, transformation of both the *mtrD* and the *mtr* promoter region yielded the donor’s azithromycin MIC of 2 μg/ml. Thus, the mechanism of resistance in mosaics is likely derived from both structural changes to *mtrD* coupled with promoter mutations that result in regulatory changes to *mtrCDE*.

The full-length *mtrD* was required for resistance in GCGS0276 transformants, indicating the role of within-gene epistasis, rather than a single point mutation, in generating azithromycin resistance. Two regions at the 5’ and 3’ ends of GCGS0276 *mtrD* together increased the MIC of 28Bl to 0.5 μg/ml (Figure 3). These two regions are part of the central pore of MtrD that stabilizes the trimeric organization of the protein (PN1) and the outer periplasmic region of the protein that may interact with MtrC to form a functional pump complex (PC2) [39]. Of note, none of the mutations observed between 28Bl and GCGS0276 have been shown to contribute to macrolide resistance in the orthologous proteins AcrB and MexB in other species, nor are they located in the direct contact site (residue 616) for macrolide recognition (e.g., [40–43]).

Within the *mtrCDE* operon, we observed local increases in linkage disequilibrium coupled with increases of rare mutations (Figure 1). These signatures could be explained by the recent acquisition of neutral diversity from closely related species, with too little evolutionary time for the combined effects of recombination and mutation to break down linkage of sites across imported DNA tracts, or the spread of these mutations to higher frequencies [44,45]. However, our experimental results confirm strong purifying selection on azithromycin plates after inheritance of partial mosaic haplotypes, suggesting that some of the linkage within *mtrCDE* observed in natural gonococcal populations may be driven by selection maintaining allelic combinations that increase resistance to azithromycin.

Overall, our results defining the role of mosaic *mtr* in azithromycin resistance affirm the importance of other *Neisseria* species as an antibiotic resistance reservoir for *N. gonorrhoeae*. Moreover, whereas mosaic *penA* genes arise from interspecies recombinations within a single gene and confer cephalosporin resistance through novel structural forms [14,46], our findings of horizontally acquired epistasitically interacting structural and regulatory variants in *mtr* point to the potential complexity by which antibiotic resistance can arise through the interactions of multiple loci. Interspecies mosaicism will be an important consideration for future development of sequence-based molecular resistance diagnostics, as markers designed to amplify gonococcal-specific sequence will overlook or incorrectly diagnose resistance phenotype. Thus, as the number of commensal neisserial genome sequences increase, analyses that map the patterns and extent of interspecies recombination may be a valuable guide in understanding pathways to resistance and in designing the appropriate diagnostic tools.

## MATERIALS AND METHODS

### Genome sequencing and population genomics

Sequencing libraries were prepared using a modification of Illumina’s Nextera XP protocol [47]. Samples were dual-indexed and pooled (n=15 per pool). Paired-end 150 bp sequencing was conducted on an Illumina MiSeq (Illumina Corp., San Diego, C.A.) platform located at the Harvard T.H. Chan School of Public Health to an average depth of 40x. Previously sequenced read libraries were obtained from the NCBI’s Short Read Archive (Project # PRJEB2090) and the European Nucleotide Archive (Project #PRJEB2999 and PRJEB7904) [7,32].

To determine the impact of interspecific recombination at *mtrRCDE*, we assessed patterns of allelic diversity across the *mtrR* transcriptional repressor and the *mtrCDE* pump compared to the rest of the genome for the 1102 GISP gonococcal isolate [7,32]. Reads were aligned to the FA1090 reference using Bowtie2 v.2.2.4 [48], and variants were called using pilon v.1.16 [49]. Vcftools v.0.1.12 [50] was used to merge resultant vcf files and calculate genome-wide values of π and Tajima’s D over 100-bp sliding windows, and r^2^ linkage by site. Gubbins v.2.2.0 [51] was used to predict regions of elevated SNP densities. For each isolate, BLASTn was used to identify the top hit and highest percent sequence identity for *mtrD* and the *mtr* promoter to all *Neisseria* within the NCBI database (*e*-value < 10^−40^).

The extent of exclusive ancestry between *N. gonorrhoeae*, *N. meningitidis, N. lactamica*, and *N. polysaccharea* was assessed using *gsi* [33] for each gene across a 25 kb window surrounding *mtrRCDE*. In brief, we downloaded *de novo* assemblies and raw sequencing reads from NCBI for *N. meningitidis* (n=431), *N. lactamica* (n=326), *N. polysaccharea* (n=37), and *N. gonorrhoeae* (n=1102; [7,32]). Raw reads were assembled with SPAdes v.3.7.0 [52], and assemblies were aligned to the *N. gonorrhoeae* FA1090 reference genome (AE004969.1) using progressiveMauve ([53]; snapshot 2015-02-13 for linux-x64), since a multi-genome alignment for all genomes was not computationally tractable. The sequences that were aligned to each gene within 25 kb of *mtrRCDE* in FA1090 were then extracted with custom Perl scripts and realigned with MAFFT v.7.309 [54]. This method of pairwise alignments of *de novo* assemblies to the FA1090 reference identifies orthologs between *N. gonorrhoeae* and the other *Neisseria* using both sequence identity and microsynteny, which is conserved in the genomic region surrounding *mtrRCDE* (Supplementary Figure 1). We then used RAxML v.8.1.4 [55] to reconstruct the phylogeny for each gene, using 50 bootstrap replicates and the GTRCAT substitution model. With these multi-species phylogenies, we calculated *gsi* with the *genealogicalSorting* R package[33,56].

### Bacterial culture conditions

*N. gonorrhoeae* isolates were provided by the CDC (Table 1). Isolates were cultured on GCB agar medium supplemented with 1% IsoVitaleX (Becton Dickinson Co., Franklin Lakes, N.J.). After inoculation, plates were incubated at 37°C in a 5% CO_2_ atmosphere incubator for 16-18 hours. Antimicrobial susceptibility testing was conducted using the agar-dilution method at a range of azithromycin concentrations from 0 to 16 μg/ml [1]. MICs were recorded after 24 hours of growth. All isolate stocks were stored at −80°C in trypticase soy broth containing 20% glycerol.

### Transformation of mosaic *mtr* alleles

Genomic DNA was extracted from isolates by lysing growth from overnight plates in TE buffer (10 mM Tris pH 8.0, 10 mM EDTA) with 0.5 mg/μl lysozyme and 3 mg/μl proteinase K (Sigma-Aldrich Corp., St. Louis, M.O.). DNA was purified using the PureLink Genomic DNA Mini Kit, treated with RNase A (Thermo Fisher Corp., Waltham, M.A.), and stored in water. Primers were designed to amplify regions of *mtrRCDE*. For primer pairs that did not amplify over a region containing a DNA uptake sequence, to enhance transformation efficiency the 12-bp AT-DUS was added to the forward primer (5’-ATGCCGTCTGAA-3’) [57]. PCR reactions were conducted in 50 μl volumes using Phusion High-Fidelity DNA Polymerase (New England Biolabs Inc., Ipswitch, M.A.) using the conditions listed in Supplementary Table 4. Amplified products were run on a 0.8% agarose gel, excised, and purified with the QIAEX II Gel Extraction Kit (Qiagen Inc., Valencia, C.A.) to remove gDNA contamination.

Transformations were conducted in GCP liquid broth (7.5 g Protease peptone #3, 0.5 g soluble starch, 2 g dibasic K_2_HPO_4_, 0.5 g monobasic KH_2_PO_4_, 2.5 g NaCl, ddH_2_O to 500 ml; Becton Dickinson Co., Franklin Lakes, N.J.) supplemented with 1% IsoVitaleX and 10 μM MgSO_4_ (Sigma-Aldrich Corp., St. Louis, M.O.). Naturally competent cells were incubated for 10 minutes with gDNA or purified PCR products to allow for DNA uptake and homologous recombination. Cells were plated on GCB with 1% IsoVitaleX and incubated for 4 hours at 37°C in a 5% CO_2_ atmosphere to allow for expression of novel alleles. Cells on expression plates were resuspended in tryptic soy broth (Becton Dickinson Co., Franklin Lakes, N.J.) and selected on 0. 38-1 μg/ml AZI GCB plates containing 1% IsoVitaleX. After 18 hours, single colonies were picked. Sanger sequencing performed using the GeneWiz sequencing service (GeneWiz Inc., Cambridge, M.A.) confirmed successful transformation.

### Transcriptome construction

Cells harvested from overnight plates were suspended in GCP supplemented with 1% IsoVitaleX and 0.042% sodium bicarbonate. Cultures were incubated at 37°C for 2 hours to midlog phase and then exposed to a sub-lethal dose of AZI (0.125 μg/ml). RNA was extracted at 0 minutes (pre-AZI) and 120 minutes (post-AZI) exposure using the Direct-Zol kit (Zymo Research, Irvine, C.A.). Transciptome libraries were prepared at the Broad Institute at the Microbial ‘Omics Core using a modified version of the RNAtag-seq protocol [58]. 500 ng of total RNA was fragmented, depleted of genomic DNA, dephosphorylated, and ligated to DNA adapters carrying 5’-AN_8_-3’ barcodes of known sequence with a 5’ phosphate and a 3’ blocking group. Barcoded RNAs were pooled and depleted of rRNA using the RiboZero rRNA depletion kit (Epicentre, Madision, W.I.). Pools of barcoded RNAs were converted to Illumina cDNA libraries in 2 main steps: 1) reverse transcription of the RNA using a primer designed to the constant region of the barcoded adaptor with addition of an adapter to the 3’ end of the cDNA by template switching using SMARTScribe (Clontech, Mountain View, C.A.) as described [59]; 2) PCR amplification using primers whose 5’ ends target the constant regions of the 3’ or 5’ adaptors and whose 3’ ends contain the full Illumina P5 or P7 sequences. cDNA libraries were sequenced on the Illumina. Nextseq 500 platform to generate 50-bp paired end reads.

Barcode sequences were removed, and reads were aligned to the FA1090 reference genome. Reads counts were assigned to genes and other genomic features using custom scripts. For the FA1090 we mapped reads to either the sense or anti-sense strand for coding domain sequences (CDSs, n=1894), tRNAs (n=55), and rRNAs (n=12). For intergenic regions (IGRs, n=1722), we mapped to each antiparallel strand. Differential expression analysis was conducted in DESeq2 v.1.10.1 [60].

## ACKNOWLEDGEMENTS

We thank David Trees and Steve Johnson for training on gonococcal transformation design and Jonathan Livny for his technical assistance with transcriptome library preparation.

